# First evaluation and detection of ivermectin resistance in gastrointestinal nematodes of sheep and goats in South Darfur, Sudan

**DOI:** 10.1101/2024.03.20.585868

**Authors:** Khalid M. Mohammedsalih, Abdoelnaim I. Y. Ibrahim, Fathel-Rahman Juma, Abdalhakaim A. H. Abdalmalaik, Ahmed Bashar, Gerald Coles, Georg von Samson-Himmelstjerna, Jürgen Krücken

**Affiliations:** Central Research Laboratory of Darfur Universities, Mousseh district, 63311 Nyala, Sudan; Faculty of Veterinary Science, University of Nyala, Mousseh district, 63311 Nyala, Sudan; Heronswey, Frog Lane, Ubley, Bristol BS40 5DU, UK; Institute for Parasitology and Tropical Veterinary Medicine, Freie Universität Berlin, Robert-von-Ostertag-Str. 7, 14163 Berlin, Germany

**Keywords:** Ivermectin resistance, *Haemonchus contortus*, sheep, goats, South Darfur, Sudan

## Abstract

In Sudan, resistance to benzimidazoles has been reported recently in cattle and goats from South Darfur. Herein, ivermectin efficacy against gastrointestinal nematodes (GINs) was evaluated in sheep and goats in three study areas in South Darfur. The faecal egg count reduction test (FECRT) was used to evaluate the efficacy of ivermectin in sheep and goats naturally infected with GINs in the region of Bulbul (goats: *n*=106), Kass (goats: *n*=40) and Nyala (Domaia (sheep: *n*=47, goats: *n*=77) and the University farm (goats: *n*=52)), using different treatment plans, and the efficacy was evaluated 12 days after treatment. Ivermectin efficacy was also evaluated in goats experimentally infected using local *Haemonchus contortus* isolates from Kass and Nyala. Nematodes surviving ivermectin treatment in goats in Bulbul and Nyala were harvested and larvae used to infect worm-free male sheep (*n*=6, ≤6 months old). Infected sheep were dosed subcutaneously with ivermectin every eight days with increasing doses from 0.2 mg/kg to 1.6 mg/kg bodyweight (bw). Reduced ivermectin efficacy was identified in sheep and goats in the four study locations. Using a paired statistic, the efficacy of a therapeutic dose in sheep was 75.6% (90% upper credible limit (UCrL): 77.5%), while twice the recommended dose led to a reduction of 92.6% (90% UCrL: 93.3%). In goats, the FECRs of a therapeutic dose were 72.9 – 95.3% (90% UCrL range: 73.6 – 95.7%) in Bulbul, Nyala Domaia, Nyala University farm and Kass. Twice the dose recommended for goats in Bulbul revealed a 90% UCrL of 87.6%. All post-treatment faecal cultures contained only *Haemonchus* spp. larvae. The experimental infection trials in sheep and goats supported our findings from field trials and calculated upper 90% CrL of below 98.9%. For the first time highly ivermectin resistant *H. contortus* populations have been identified in sheep and goats in Sudan, and resistance was experimentally confirmed.

## Introduction

Parasitic worms are an important public health and veterinary problem worldwide. In tropical regions, including Sudan, helminths, such as nematodes, are able to cause significant morbidity and mortality in their hosts [1, 2]. In livestock, gastrointestinal nematodes (GINs) are causing significant losses in production. The strongest pathogenic effects in ruminants are due to infection with *Haemonchus* spp., haematophagous parasites in the abomasum [3]. *Haemonchus contortus* is an economically important parasite of small ruminants, and it causes serious problems in both extensive and intensive small ruminant farming systems in warm to moderate climatic regions. Due to blood losses during feeding and continuous bleeding at previous attachment sites, anaemia, hypoproteinaemia, oedema and loss of animal live-weight are frequently observed clinical signs [4]. Infected animals also tend to have a reduced digestive capacity which affects the uptake of nitrogen, organic matter and energy, and death may occur in heavily infected young animals [3]. Other GIN genera are also described as pathogenic and causing significant production losses in sheep and goats, including the genera *Teladorsagia*, *Cooperia*, *Trichostrongylus*, *Chabertia*, *Oesophagostomum* and *Nematodirus* [5]. Importantly, co-infections with two or more of these species regularly occur in the field with complicating the clinical consequences and potentially also the treatment requirements [6].

Currently, application of anthelmintic drugs remains a core control strategy against GIN infections in humans and animals. Macrocyclic lactones, benzimidazoles and imidazothiazoles are the major anthelmintic drug classes marketed in the veterinary pharmacies over the last six to four decades for treatment of the infected animals [7]. Ivermectin, belonging to the macrocyclic lactones, has been marketed since the 1990s as a highly effective and broad-spectrum anthelmintic drug also covering many ectoparasites such as ticks and lice. Besides its broad-spectrum efficacy, ivermectin has a wide safety margin making it the drug of choice against parasitic nematodes and arthropods in domestic animals [8]. The widespread use of ivermectin in the last decades has led to the evolution of resistance, which has been initially reported from countries with intensive ruminant farming such as Australia, New Zealand and South Africa and is now also found widespread in South America, the United Kingdom and the European Union [9–12]. In Sub-Saharan Africa (excluding South Africa with its intensive farming system), resistance to ivermectin has been reported for subsistence farming systems in sheep, goats and cattle in many regions, including Ethiopia [13], Kenya [14], Nigeria [15] and Uganda [16]. To the best of our knowledge respective data are not yet available for Sudan.

Sheep and goats show differences in bioavailability of anthelmintics including ivermectin. The bioavailability of ivermectin is remarkably different in goats compared to sheep, therefore goats require (1.5 to) 2-fold the sheep recommended dose to achieve the same pharmacological effect [17–19]. In Sudan (Mohammedsalih, personal observation) as in many Sub-Saharan countries [14, 15, 20], the package labels for ivermectin are identical for sheep and goats stating the use of 0.2 mg/kg body weight (bw) for goats which corresponds to a constant under-dosing and plays a central role in the evolution of high level ivermectin resistance in goats [18].

In recent years, efforts have focussed on understanding the situation of anthelmintic resistance in Sudan. Resistance to benzimidazoles has been studied in cattle and goats in South Darfur, with detection of widespread benzimidazole-resistant *H. contortus* populations in small ruminants but to some extend also in cattle [2, 21]. The objective of this study was to investigate the efficacy of ivermectin in sheep and goats naturally infected with GINs in South Darfur. Moreover, ivermectin efficacy was evaluated in goats experimentally infected with *H. contortus* isolated from a local abattoir and sheep experimentally infected by nematodes surviving ivermectin treatment in goats.

## Materials and methods

### Study location and design

The study was conducted in South Darfur (11.30°N 24.40°E), southwestern Sudan. Three areas were included; Bulbul (12.43°N 24.29°E), Kass (12.50°N 24.28°E) and Nyala (12.05°N 24.88°E). Two subareas were represented in Nyala; Domaia village, in western direction, and the University farm in the southeast direction of Nyala.

South Darfur is in a savannah zone and mostly covered by a clay-sandy soil, and the abo-asabei (Egyptian crowfoot grass, *Dactyloctenium aegyptium*) is the dominant grass. The state has a long dry season with a single rainy season from June to November with a precipitation range between 377 – 546 mm/month in the rainy season. The average minimum and maximum temperatures are 24.7 °C and 37.6 °C, and the relative humidity is in the range of 28.3 – 56.7%. Subsistence farming is the main practiced system in the three selected areas, where livestock rearing is integrated with crop production [22, 23]. The University farm is an educational farm at the Faculty of Veterinary Science, University of Nyala, Nyala, Sudan, keeping and breeding local sheep, goat and cattle breeds. The practiced farm management is a semi-intensive system. During the daylight, animals have unrestricted access to pasture inside the University campus (16 km^2^), which is mostly dominated by abo-asabei grass. In the dry seasons, the animals feed concentrates as a supplementation.

Ivermectin efficacy was evaluated in local sheep (*Ovis aries*) and goat (*Capra hircus*) breeds, desert sheep and goats, naturally infected with GINs in regions of Bulbul, Kass and Nyala based on the faecal egg count reduction test (FECRT) during the rainy season (June – November) in the years 2019 – 2022. The efficacy against *H. contortus* was also studied in goats experimentally infected with *H. contortus* isolated from goats in the Kass and Nyala abattoirs. Furthermore, resistance to ivermectin was confirmed in sheep experimentally infected with third stage larvae (L3) from nematodes surviving ivermectin treatment in goats

Although the revised World Association for the Advancement of Veterinary Parasitology (WAAVP) guidelines were not published at the time of the study, the study design was completely compliant with the revised guideline regarding the recommended number of included animals and the total number of observed nematode eggs. The nematode eggs per gram (EPG) of faeces of the treatment group animals was also determined before treatment, which allowed the calculation of the FECR based on paired data. The total number of strongyle eggs counted before treatment was always above 200, which was recommended by Kaplan [24] and well within or even above the range recommended for ruminants (Box3 in Kaplan et al. [25]). This was achieved by including only animals with an EPG above 150 and group sizes between 15 and 78 animals. In South Darfur, the owners of the sheep and goats usually practice subsistence farming, only owning a very small number of animals, frequently sharing the pastures with other owners, where available. Therefore, ivermectin efficacy was herein studied at a regional (as defined above) and not a farm level. When selected animals on individual farms were less than 15, sheep and goats from two or three neighbouring farms were considered to be one treatment group to achieve at least 15 animals per treatment group.

### Ivermectin treatment in sheep and goats naturally infected with gastrointestinal nematodes

Ivermectin efficacy was studied on 46 farms (median 8 animals, range 4 – 34) in three different study areas in South Darfur. In any study area, all farms were visited and included if they had at least four sheep and/or goats which were not treated with any anthelmintic for at least 30 days. Farms were distributed as follows: Bulbul (*n*=18), Kass (*n*=10) and Nyala Domaia (*n*=17). Since the total number of sheep in each farm in Bulbul, Kass and at the Nyala University farm was between 0 to 4 heads/farm (median 0 sheep), while the total number in Nyala Domaia was much higher (4 – 25 heads/farm), the efficacy in sheep was studied only in the latter area. The University farm had a total of 98 goats. On each farm, in the initial screening untreated male and female sheep and goats of different age groups (young; <1 year, adult; >1years, based on dentition [26]) were examined for the presence of GINs infection using the Mini-FLOTAC device (multiplication factor of 10). Animals with more than 150 EPG for strongyle nematodes [25] were labelled to be included in the trials. A total of 47 sheep and 275 goats, respectively, were selected to be included in FECRTs. From the selected animals, 37 (78.7%) sheep and 230 (83.6%) goats received ivermectin and were available at the trial end (Table 1).

**Table 1.** Sheep and goats used to assess the efficacy of ivermectin in three different study areas in South Darfur, Sudan.

| Study animal | Factor | Total | Sex |  | Age |  | Study area |  |  |  |
| --- | --- | --- | --- | --- | --- | --- | --- | --- | --- | --- |
|  |  |  | male | female | young | adult | Bulbul | Kass | Nyala Domaia | Nyala University farm |
| Sheep | All tested sheep | 68 | 10 | 58 | 32 | 36 | – | – | 68 | – |
|  | No. (%) of sheep shedding strongyle eggs | 54 (79.4) | 7 (70.0) | 47 (81.0) | 26 (81.3) | 28 (77.8) | – | – | 54 (79.4) | – |
|  | No. (%) of sheep shedding >150 strongyle eggs per gram faeces | 50 (73.5) | 7 (70.0) | 43 (74.1) | 24 (75.0) | 26 (72.2) | – | – | 50 (73.5) | – |
|  | No. (%) of selected sheep that were treated and available at the trial end | 37 (54.4) | 6 (60.0) | 31 (53.5) | 14 (43.8) | 23 (63.9) | – | – | 37 (54.4) | – |
| Goats | All tested goats | 395 | 45 | 350 | 117 | 278 | 120 | 80 | 115 | 80 |
|  | No. (%) of goats shedding strongyle eggs | 315 (79.8) | 28 (62.2) | 287 (82.0) | 65 (55.6) | 250 (89.9) | 111 (92.5) | 53 (66.3) | 90 (78.3) | 61 (76.3) |
|  | No. (%) of goats shedding >150 strongyle eggs per gram faeces | 286 (72.4) | 25 (55.6) | 261 (74.6) | 52 (44.4) | 234 (84.2) | 110 (91.7) | 44 (55.0) | 80 (69.6) | 52 (65.0) |
|  | No. (%) of selected goats that were treated and available at the trial end | 230 (58.2) | 21 (46.7) | 209 (59.7) | 52 (44.4) | 178 (64.0) | 96 (80.0) | 30 (37.5) | 62 (53.9) | 42 (52.5) |

The body weight of the animals was determined before ivermectin treatment using the measurement of heart girth and body length [27] and calculated according to the formula:

> Heart girth × heart girth × body length / 600 = animal weight in kilograms

Ivermectin (IVOMEC^®^ 10 mg/ml solution for injection (batch No. BK253), MERIAL GmbH, Hallbergmoos, Germany; Virbamec^®^ 1.0% w/v solution for injection (batch No. 4ZJZ), Warnham, United Kingdom; and Intermectin 0.08% drench (batch No. 362874), Interchemie, Venray, The Netherlands) was imported from Europe and used either as subcutaneous (SC) injection or orally at different dose regimens. Sheep (*n*=37) were divided into two treatment groups, the first treatment group (*n*=22) received ivermectin (IVOMEC^®^) SC at a therapeutic dose (0.2 mg/kg bw) [28]), while the second group (*n*=15) was treated with twice the recommended dose (0.4 mg/kg bw).

In goats, ivermectin efficacy was evaluated based on different dose regimens in the three selected areas. The recommended therapeutic dose for goats (0.4 mg/kg bw [28]) was administered (IVOMEC^®^ or Virbamec^®^) SC to 212 male and female goats of different age groups that were distributed as follows: Bulbul (*n*=78), Kass (*n*=30), Nyala Domaia (*n*=62) and Nyala University farm (*n*=42). Twice the recommended goat dose (0.8 mg/kg bw) was tested in 18 previously untreated goats in Bulbul. Goats (*n*=63) treated with 0.4 mg/kg dose in Bulbul (*n*=33) and Nyala University farm (*n*=30) that continued shedding >150 strongyle EPG were retreated on day 12 with 0.8 mg/kg bw dose. IVOMEC^®^ was SC injected in goats in Bulbul, Kass and the Nyala University farm, while Virbamec^®^ brand was only administered to goats from Nyala Domaia. In all of these trials, faecal samples were collected before treatment (day 0) and on day 12 after ivermectin administration. Study animals stayed on their farms throughout the experiment and after testing was finished.

### Ivermectin treatment in goats experimentally infected with Haemonchus contortus

Ivermectin efficacy was further studied in two separate experimental trials using two different *H. contortus* isolates from Nyala and Kass. Experimental trials were conducted at the premises of the Faculty of Veterinary Science, University of Nyala. Experimental infections were performed as described by Mohammedsalih et al. [2]. Briefly, at each abattoir (Kass and Nyala) 50 abomasa from naturally infected goats were collected. Mature gravid female *H. contortus* were isolated from these abomasa, crushed, pooled separately for each abattoir and cultured in heat-treated bovine faeces. The culture medium was prepared by collecting faeces from two worm-free calves, tested using the Mini-FLOTAC device, dried and heated to 70 °C for 2 h. After 10 days of incubation at 22 – 27 °C, the Baermann method was used to harvest infective L3 from faecal cultures and the number of L3 was determined [29]. The experimental design and treatment plans of these trials are summarized elsewhere in Table 2. For the two experimental infection trials, 30 clinically healthy male goats (Kass experiment; *n*=10, Nyala experiment; *n*=20) of 3 – 6 months age and apparently negative for faecal egg shedding, based on Mini-FLOTAC technique, were weighed (6 – 18 kg) and all purchased goats (*n*=30) were prophylactically treated with a single dose of levamisole SC (10 mg/kg bw, levamisole 100 mg/kg Inj.-Lsg., cp-pharma (batch No: 3893), Burgdorf, Germany). After 21 days, faecal samples were examined and negative animals were inoculated orally with 150 L3/kg bw [30]. Starting 10 days after inoculation, faecal samples were collected daily and examined for the presence of strongyle eggs. At day 25 post infection, male goats of each of the two separate experiments, infected with *H. contortus* from either Kass or Nyala, were divided into three treatment groups. The trial in Kass had a single treated group of 10 animals, while two treatment groups (10 male goats each) were included in the Nyala experiment to compare different administration routes. Since oral administration of ivermectin is commonly practiced by farmers in South Darfur (Mohammedsalih, personal observation), the first treatment group infected with *H. contortus* isolates from Nyala received an oral dose of 0.4 mg/kg bw ivermectin (Intermectin 0.08% drench), while the second treatment group and the treated group infected by *H. contortus* isolates from Kass were treated with the same dose (IVOMEC^®^) using the SC route (Table 2). Throughout the study period, goats were fed dry hay with free access to drinking water, and care was taken to avoid any contamination of pens with nematode larvae from outside. Faecal egg counts were determined, for each group, before treatment (day 0), then eight, ten, twelve and 14 days after administration of ivermectin.

**Table 2.**
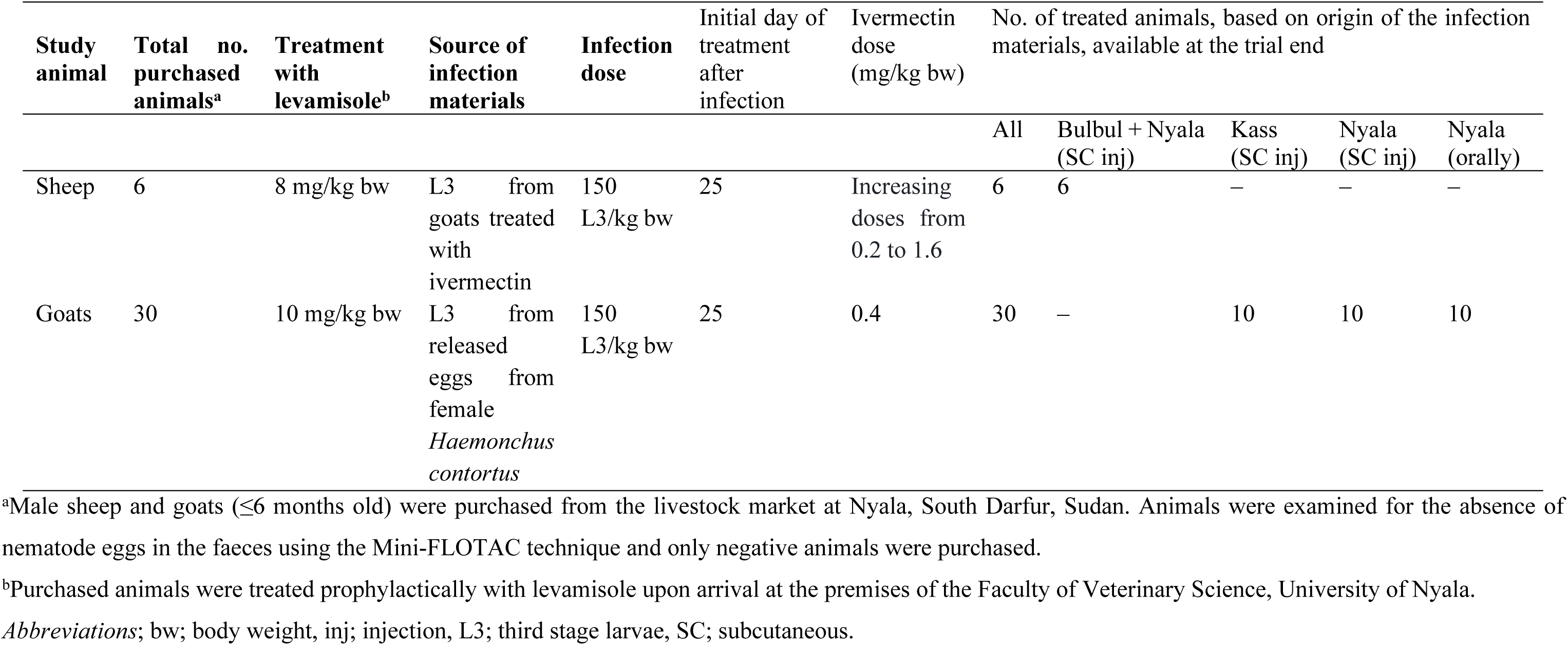
Study design of experimental infections of sheep and goats.

| Study animal | Total no. purchased animals<sup>a</sup> | Treatment with levamisole<sup>b</sup> | Source of infection materials | Infection dose | Initial day of treatment after infection | Ivermectin dose (mg/kg bw) | No. of treated animals, based on origin of the infection materials, available at the trial end |  |  |  |  |
| --- | --- | --- | --- | --- | --- | --- | --- | --- | --- | --- | --- |
|  |  |  |  |  |  |  | All | Bulbul + Nyala (SC inj) | Kass (SC inj) | Nyala (SC inj) | Nyala (orally) |
| Sheep | 6 | 8 mg/kg bw | L3 from goats treated with ivermectin | 150 L3/kg bw | 25 | Increasing doses from 0.2 to 1.6 | 6 | 6 | – | – | – |
| Goats | 30 | 10 mg/kg bw | L3 from released eggs from female *Haemonchus contortus* | 150 L3/kg bw | 25 | 0.4 | 30 | – | 10 | 10 | 10 |

<sup>a</sup>Male sheep and goats ( $\leq 6$ months old) were purchased from the livestock market at Nyala, South Darfur, Sudan. Animals were examined for the absence of nematode eggs in the faeces using the Mini-FLOTAC technique and only negative animals were purchased.
<sup>b</sup>Purchased animals were treated prophylactically with levamisole upon arrival at the premises of the Faculty of Veterinary Science, University of Nyala.
*Abbreviations*; bw; body weight, inj; injection, L3; third stage larvae, SC; subcutaneous.

### Ivermectin treatment in sheep experimentally infected with nematodes surviving ivermectin treatment in goats

This experiment aimed to confirm presence of ivermectin resistance in parasitic nematodes surviving ivermectin treatments in goats in Bulbul and Nyala University farm. For this purpose, sheep were experimentally infected with L3 obtained from such treated goats, as summarized in Table 2. Goats that continued shedding nematode eggs after treatment with 0.8 mg/kg bw dose in Bulbul and those retreated with 0.8 mg/kg bw ivermectin on day 12 after the first treatment with 0.4 mg/kg bw in Bulbul and Nyala University farm were used to collect infection material, which was pooled from these groups. Faecal samples were collected on day 12 post treatment, pooled, cultured, incubated and L3 were harvested using the Baermann method [29]. Six clinically healthy male sheep, 3 – 6 months old, tested negative for infection with GINs (based on the Mini-FLOTAC technique) were located at the premises of the Faculty of Veterinary Science, University of Nyala. Study sheep had free access to drinking water and they were fed dry hay during the study period. The pen was monitored to avoid any contamination with nematode larvae from outside. Three weeks prior to infection, study sheep were weighed (7 – 20 kg) and treated subcutaneously with 8 mg levamisole/kg bw. After three weeks, each sheep was tested for the absence of nematode eggs before receiving an oral dose of 150 L3/kg bw [30]. From day 10 after inoculation, faecal samples were collected every 24 hours until detection of nematode eggs. At day 25 post infection, sheep were dosed SC with ivermectin (IVOMEC^®^) every eight days with increasing doses starting from the dose recommended for sheep (0.2 mg/kg bw) followed by the ivermectin dose usually recommended for goats (0.4 mg/kg bw), then 0.8 mg/kg bw and 1.6 mg/kg bw. Faecal egg counts were conducted before treatment (day 0), then every eight days up to day 28 after SC injection of the highest dose (1.6 mg/kg bw). Faecal samples were collected on day 8 of 0.2 mg/kg bw dose administration (actually, day 33 of infection), then 16 days from starting the first dose of ivermectin administration (actually, day 8 from administration of the second dose, 0.4 mg/kg bw), day 24 (actually, day 8 from administration of the third dose, 0.8 mg/kg bw) and day 32 (actually, day 8 from administration of the fourth dose, 1.6 mg/kg bw). Ivermectin dosing was stopped at 1.6 mg/kg bw, however the collection of faecal samples was continued to days 39, 46 and 53 after the first day of ivermectin treatment. After the end of the experiment, study sheep were slaughtered at Nyala abattoir according to the halal method [31]. The four stomaches and the intestine were isolated and inspected for the presence of immature and adult GINs.

### Faecal sample analyses

#### Faecal egg counts

Individual faecal samples were collected directly from the rectum of sheep and goats in plastic bags, labelled and stored at 4 °C for a maximum of 24 h before egg counting. Faecal samples from Bulbul and Nyala trials were analysed at the Laboratory of Parasitology, Faculty of Veterinary Science, University of Nyala, while those collected from Kass were analysed directly in the field. Faecal samples were prepared as explained by Mohammedsalih et al. [2] using the Mini-FLOTAC device [32] and nematode eggs were differentiated and counted using standard keys [29]. The EPG was calculated by multiplying the number of eggs observed under the light microscope (Olympus, Philippines) with ten.

#### Faecal cultures

Faecal samples were collected from sheep and goats naturally infected with GINs, that were selected to evaluate the efficacy of ivermectin on day 0 (before treatment) and 12 days after administration of the drug, and pooled on the farm level for each time point. Faeces were mixed with wood shaving and incubated in labelled plastic jars at 22 – 27 °C with daily moistening using distilled water for 10 days [29]. Third stage larvae were purified with the Baermann method. For larval identification, the first 100 L3, or if less than 100 all L3, were identified under a light microscope, and assigned to the genera *Haemonchus*, *Trichostrongylus* and the group *Oesophagostomum/Chabertia* using standard differentiation keys [29, 33].

### Statistical analyses

The results of the FECRT were evaluated according to the guideline of the WAAVP. Since this guideline has been updated only recently [25], the effects of this revision were investigated by comparing interpretations based on the original and the revised guideline. Since the study was designed in 2019, when neither the revised WAAVP guideline [25] nor the suggestions by Kaplan [24], which already included many of the study design principles proposed in the revised guideline, were available, the study design was still largely according to the rules of the original guideline [34]. However, since this study was completely compliant with the revised guideline regarding the recommended number of included animals and the total number of observed eggs, the analysis of the data and interpretation of the results was continued according to the revised guideline [25]. In addition, a paired data analysis of pre and post treatment data of all field trial data of different trials was further interpreted according to the original guideline to compare between the two guidelines.

The faecal egg count reductions (FECRs) were calculated to assess the efficacy of ivermectin in sheep and goats that were naturally infected with GINs at 12 days after treatment. For the experimental infection trials in sheep and goats, the efficacy was evaluated at different time points. The paired FECRs with 90% and 95% credible intervals (CrIs) were calculated in R 4.1.1 using the eggCounts Bayesian hierarchical model version 2.3-2 without zero-inflation option and without individual efficacy [35]. The package coda 0.19-4 was used to extract 90% CrI as 90% highest posterior density intervals from the eggCounts results. According to the research protocol for ruminants in the revised WAAVP guideline [25], the expected efficacy was set to 99% and the lower efficacy target to 95%. Thereafter, the efficacy was classified exclusively based on the 90% CrIs. A parasite community was classified as ivermectin susceptible when the lower 90% credible limit (CrL) was equal to or greater than the lower efficacy target of 95%. Resistance was considered when the upper 90% CrL was less than the expected efficacy of 99%. The new sub-classification, low resistance, was assigned when the upper 90% CrL was below 99% and the lower 90% CrL was equal to or greater than 95%. If neither of these conditions were met, i.e. the lower 90% CrL was lower than 95% and the upper CrL was above 99%, the FECRT result was considered to be inconclusive. For comparison, the results were also interpreted according to the original guideline [34] and a community was considered to be resistant when the estimate of the FECR was below 95% and the lower 95% CrL was less than 90%. Parasitic nematodes were considered to be susceptible when the FECR was above 95% and its lower 95% CrL was at least 90%. Otherwise, when the FECR estimate was below 95% and the lower 95% CrL was at least 90%, the FECRT was considered to be inconclusive, or “suspected resistance”.

In addition to the above mentioned methods used for calculating and classifying resistance or susceptibility, the Bayesian-Frequentist Hybrid Inference method; an open-source software using the R package bayescount version 1.1.0 was accessed through a web interface (https://www.fecrt.com) and used to determine the efficacy status of a parasitic nematode community [36] by calculating the 90% CrI based on the delta method [37]. Since the method does not provide an estimate for the FECR, the percent reduction of mean FEC data of pre and post-treatment samples were used as a simple estimate for the FECR. The results of the bayescount package were only interpreted using the revised guideline since the method does not by itself calculate an estimate of the FECR, which is required for interpretation according to the original guideline. The widths of the CrI were compared between eggCounts and bayescount methods using a Wilcoxon matched-pairs signed rank test in GraphPad version 8.0.2. Agreement of assignments between the statistical methods or the two guideline was analysed using Cohen’s κ statistics calculated using the CohenKappa() function from DescTools 0.99.48.

In order to evaluate the effects of study regions, host species, age, sex, the ivermectin dose and the route of administration on the efficacy of ivermectin in sheep and goats naturally infected with GINs logistic regression models were calculated. For this purpose, individual faecal egg count reduction (FECRi) was calculated for each of the study animals using the following formula [38, 39]:

> FECRi (%) = 100 × [1 – (T_i_/T_0_)]

In this formula, T_i_ represents the EPG after treatment and T_0_ represents the EPG before treatment (day 0). These FECRi values were then transformed to binomial variables (FECRi99 and FECRi95) using two different thresholds for full susceptibility, i.e. setting all FECRi ≥99% or all FECRi ≥95% to 1 and all values below the threshold to 0. Thereafter, the obtained data were used to calculate the logistic regressions using the explanatory variables; study regions (Kass vs. Bulbul, Nyala Domaia and Nyala University farm), host species (sheep vs. goats), sex (male vs. female), age (young vs. adult, based on dentition [26]) and the administered ivermectin dose (the therapeutic dose vs. twice the recommended dose). The models were calculated using the glm() function in R. Explanatory variables were stepwise eliminated to minimise the Akaike Information Criterion (AIC) of the model using the drop1() function. The 95% confidence intervals (CIs) were calculated using confint() and odds ratios were obtained by applying exp() to the estimates and confidence limits (CLs). The RsqGLM() function from the modEvA package 3.11 was used to calculate pseudo R^2^ values according to Nagelkerke and Tjur. A value of p < 0.05 was considered statistically significant.

### Ethics statement

Ethical approval to conduct the trials of this study was obtained from Research and Ethics Committee at the Faculty of Veterinary Science, University of Nyala, Sudan (Ref. UN/FVS/1/34). For the field trials, informed verbal consent was obtained from the farmers. Verbal consent was chosen since a large proportion of the local rural population are illiterate. This procedure was approved by the Ethics Committee.

## Results

### Ivermectin treatment in sheep and goats naturally infected with gastrointestinal nematodes

In South Darfur, a total of nine FECRTs for strongyle nematodes was conducted to evaluate the efficacy of ivermectin in sheep (number of tests = 2) and goats (number of tests = 7) naturally infected with strongyle nematodes in the four different study locations (Table 3). All raw data for these samples can be found in S1 Data. The results of the FECRT were initially evaluated using the eggCounts package and the revised guideline [25]. The tests in sheep used the recommended dose and twice the recommended dose and the strongyle communities in Nyala (Domaia) were assigned to a resistance status independently of the applied dosage (Table 3). However, when comparing the 95% CrIs, which are not overlapping, the FECR was at least significantly higher with twice the recommended sheep dose of ivermectin compared to the recommended dose (p < 0.05) (Fig 1).

**Fig 1.**
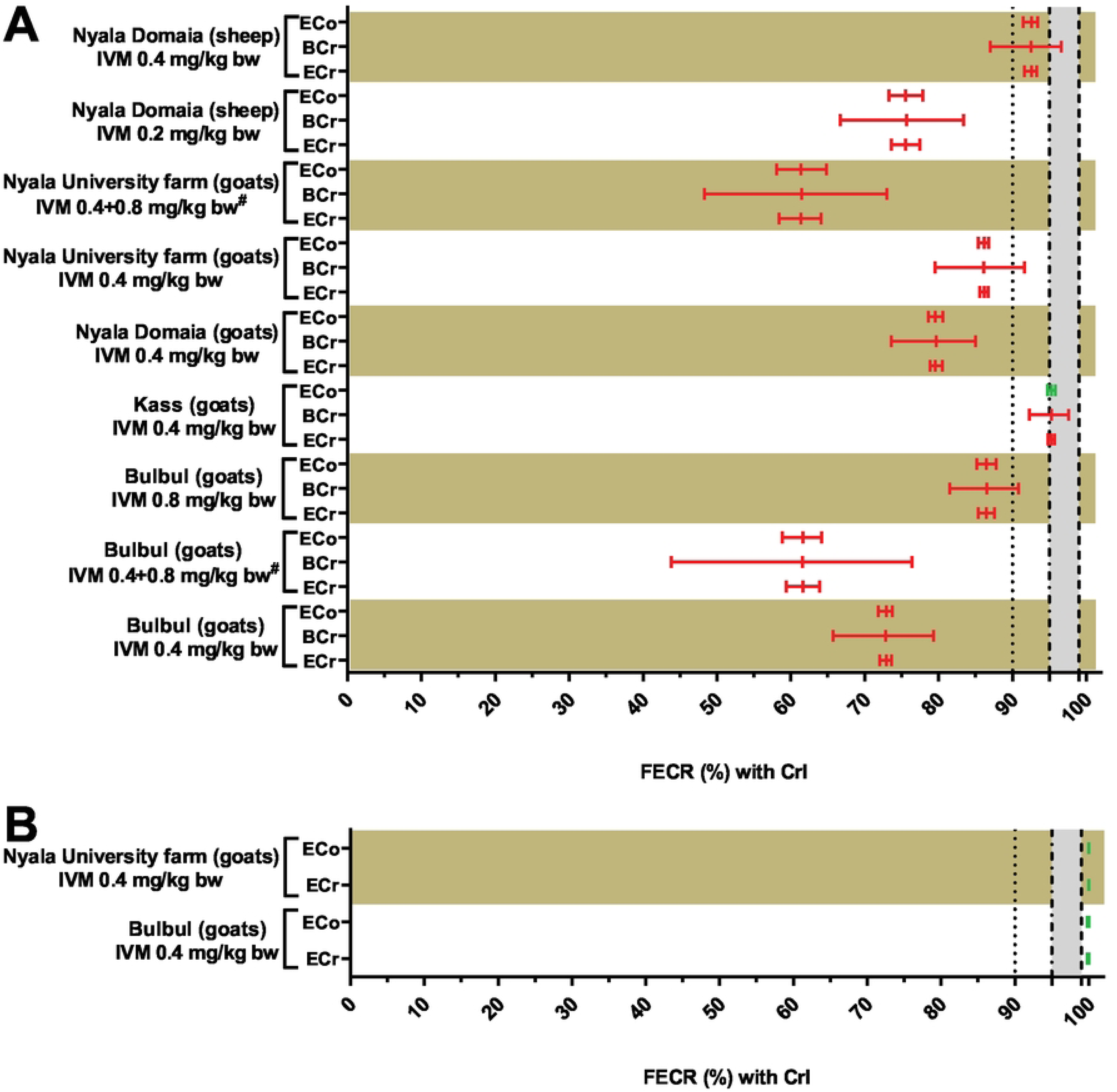
Comparisons between the faecal egg count reductions (FECRs) with both 90% and 95% credible intervals (CrIs) in sheep and goats naturally infected with gastrointestinal nematodes (strongyles (A) and *Strongyloides papillosus* (B)) after subcutaneous administration of ivermectin (IVM) at different doses to the treated groups in three different study areas in South Darfur, Sudan, using different statistical analyses methods. ECr: paired faecal egg count reductions (FECRs) and 90% credible intervals (CrIs) were calculated in R 4.1.1 using the eggCounts Bayesian hierarchical model version 2.3-2 without zero-inflation option and without individual efficacy. BCr: the 90% CrIs of the FECR for paired data were calculated using the Bayesian-Frequentist Hybrid Inference method through an open-source webpage using the R package bayescount version 1.1.0 (https://www.fecrt.com). The average reduction in FECs after treatment was calculated separately. ECo; the paired FECRs and 95% CrIs were calculated in R 4.1.1 using the eggCounts Bayesian hierarchical model version 2.3-2 without zero-inflation option and without individual efficacy. Results for ECr and BCr were assigned to the status resistance (red colour), low resistance (orange), susceptible (green) or inconclusive (black) as recommended in the revised guideline [25]. Results of the ECo were assigned resistance (red colour), susceptible (green) or inconclusive (black) as in the original guideline [34]. Calculation of CrI using the BCr was not possible for *S. papillosus* data since the CrIs are not calculable if all post-treatment data are zero. The grey zone indicates the range between the lower efficacy target of 95% and the expected efficacy of 99%. ^#^Retreated goats were initially treated with 0.4 mg/kg ivermectin (day 0) and received a twice the recommended dose of ivermectin (0.8 mg/kg body weight) on day 12.

**Table 3.**
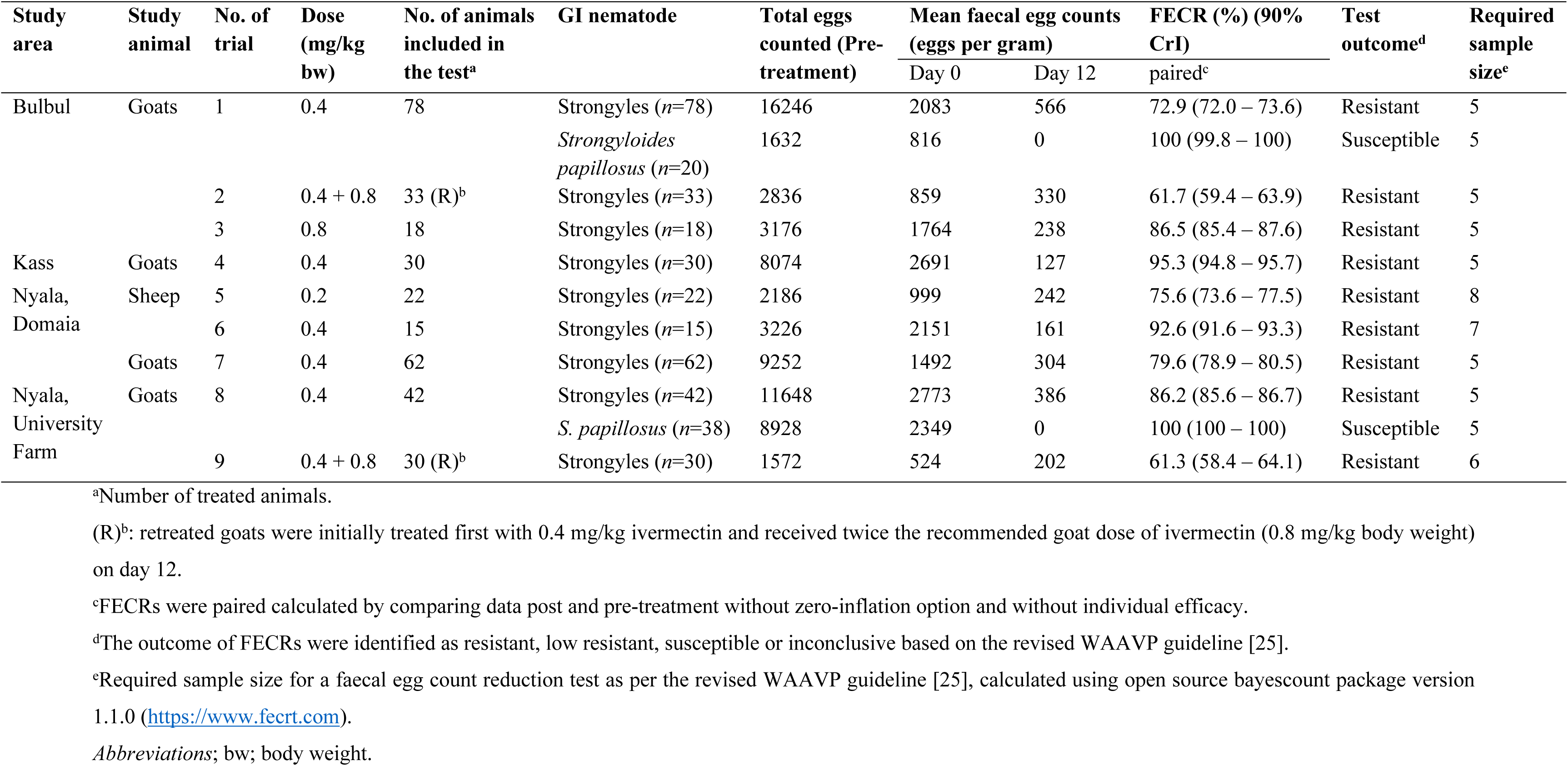
Results of the faecal egg count reduction (FECR) (and 90% credible interval (CrI)) with sheep and goats naturally infected with gastrointestinal nematodes before and after subcutaneous administration of ivermectin at different doses to the treated groups at three different study areas in South Darfur, Sudan.

In goats, all tests using either the recommended goat dose of 0.4 mg/kg bw or twice the recommended goat dose (0.8 mg/kg bw) also identified the strongyle communities as resistant to ivermectin (Table 3). In Bulbul, where both doses were tested in parallel, the FECR achieved with the higher dose was significantly higher (p < 0.05) than with the lower dose (Fig 1).

In the third treatment plan, goats in Bulbul and in the University Farm in Nyala were treated first with the recommended dose and animals that still had an EPG above 150 at day 12 were retreated with twice the recommended goat dose on day 12 after the first treatment. This second treatment led to the lowest FECR values that were observed in this series of tests (Table 3).

In two of the tests in goats, the EPG for *Strongyloides papillosus* was high enough to also calculate a FECR for this species using the recommended dose of 0.4 mg/kg ivermectin. This species was fully susceptible and no *S. papillosus* eggs were found post treatment (Table 3). Although *Strongyloides* spp. are not mentioned in the revised guideline for the FECRT [25] and thus neither an expected efficacy nor a lower target efficacy has been proposed, the observed efficacy was significantly higher (p < 0.05) than 99% and thus this parasite was considered to be fully susceptible for ivermectin.

In coprocultures, sheep and goats showed mixed infections when the faecal cultures of strongyle L3 were examined before treatment, including *Haemonchus* spp., *Trichostrongylus* spp. and Chabertiidae (*Oesophagostomum* spp./*Chabertia* spp.) with *Haemonchus* spp. showing the highest percentage (71 – 92%) (Table 4). For all treatment groups, larval cultures contained only *Haemonchus* spp. in post-treatment samples.

**Table 4.**
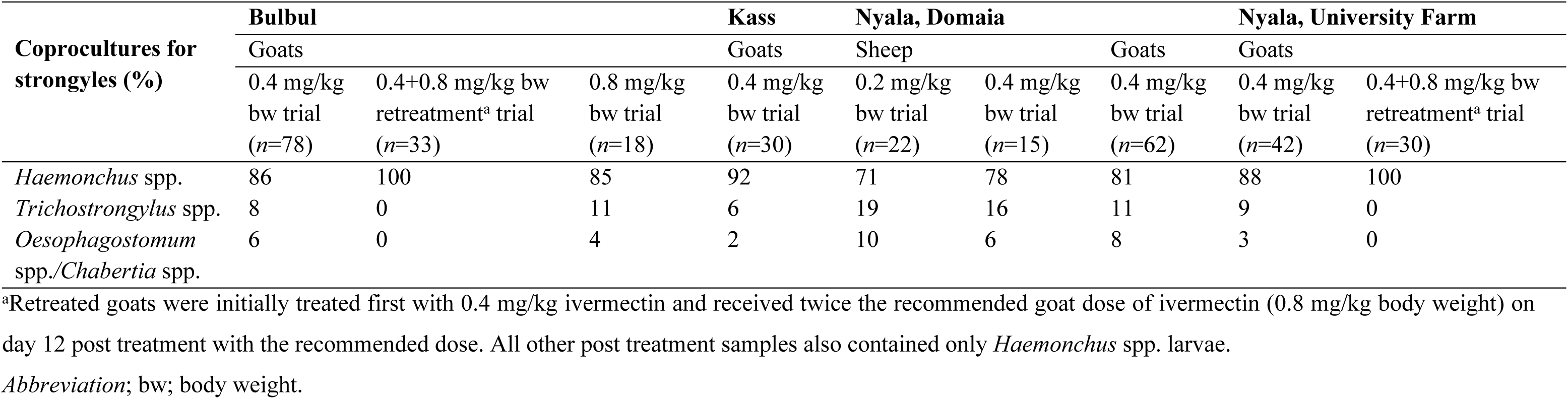
Coprocultures of gastrointestinal helminths in the faeces of naturally infected sheep and goats before treatment (day 0) at three different study areas in South Darfur, Sudan.

### Classification of ivermectin efficacy using the original and revised (research) WAAVP criteria and comparison of the statistical packages eggCounts and bayescount

All of the eleven FECRTs performed in sheep and goats naturally infected with strongyle nematodes or *S. papillosus* were evaluated based on three different methods (Fig 1): (i) eggCounts with 90% CrI using the revised guideline, (ii) bayescount with 90% CrI or the Beta Negative Binomial (BNB) method using the revised guideline and (iii) eggCounts with 95% CrI and the estimate of the FECR according to the original guideline. Since FECs of all samples were zero for *S. papillosus*, bayescount was not able to calculate any 90% CrIs for *S. papillosus* using the delta method and chose the BNB method instead, which does not provide a CrI. The comparison of eggCounts and bayescount according to the revised guideline was done for strongyles as well as *S. papillosus*. However, for comparison of the 90% CrIs only the strongyle data were included. These data showed that 90% CrI calculated with bayescount were significantly larger for all strongyle data sets compared to eggCounts (p = 0.0039, Wilcoxon matched-pair signed rank test) (Fig 2). Despite this clear difference in CrI width, both statistical methods classified all FECRT for strongyles as resistant and both FECRT for *S. papillosus* as susceptible. This complete agreement led to a Cohen’s κ value of 1. In contrast, comparison of the interpretation of the revised and the original guideline based on calculations using eggCounts revealed small differences. For one data set, strongyle parasites in goats from Kass treated with the recommended goats of 0.4 mg/kg bw, the revised guideline assigned a resistant status while the original guideline assigned it to a susceptible status (Fig 1) leading to a Cohen’s κ value of 0.74.

**Fig 2.**
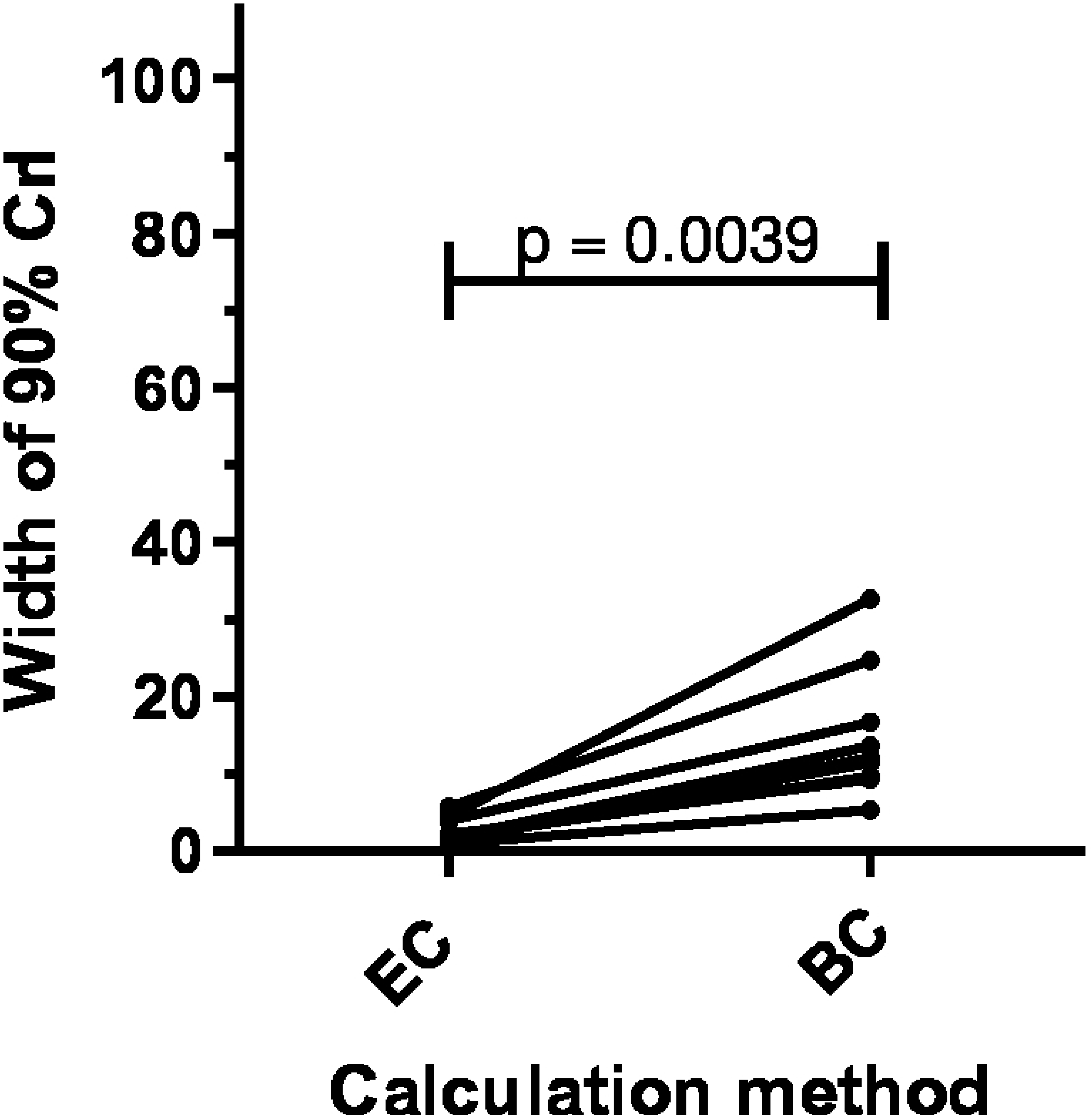
Paired values for the width of the 90% credible interval (CrI) for the eggCounts (EC) and the bayescount delta method (BC). According to the Wilcoxon matched-pairs signed rank test, the width of the 90% CrI was significantly smaller when calculated according to eggCounts in comparison to the bayescount delta method.

### Factors associated with ivermectin efficacy in individual sheep and goats

For each individual animal, the treatment was classified as efficient (1) or not efficient (0) if the FECRi was equal to or above 95% (FECRi95) or 99% (FECRi99). The explanatory variables study region, host species, age, sex, ivermectin dose and route of administration were analysed in small ruminants naturally infected with strongyle nematodes. Explanatory variables were stepwise eliminated to improve the model AIC and the final model included only the region, which was significantly (p < 0.05) associated with increased odds of poor ivermectin performance using the variables FECRi95 and FECRi99 (Table 5).

**Table 5.** Final logistic regression model to identify variables with significant effect on the odds (95% confidence interval (CI)) of goats naturally infected with gastrointestinal nematodes in three different study areas in South Darfur, Sudan, to treatment with ivermectin.

|  | 95% efficacy |  |  |  |  | 99% efficacy |  |  |  |  |
| --- | --- | --- | --- | --- | --- | --- | --- | --- | --- | --- |
|  | Estimate | Standard error | p- value | Odd ratio | 95% CI | Estimate | Standard error | p- value | Odd ratio | 95% CI |
| **Intercept** | 0.406 | 0.373 | 0.277 | 1.500 | 0.730 – 3.197 | -0.268 | 0.368 | 0.467 | 0.765 | 0.364 – 1.568 |
| **Study area: Kass vs.** |  |  |  |  |  |  |  |  |  |  |
| Bulbul | -2.674 | 0.511 | <0.0001 | 0.069 | 0.024 – 0.182 | -3.582 | 0.804 | <0.0001 | 0.028 | 0.004 – 0.112 |
| Nyala Domaia | -1.548 | 0.476 | 0.0012 | 0.213 | 0.081 – 0.532 | -1.380 | 0.505 | 0.0063 | 0.252 | 0.091 – 0.670 |
| Nyala University farm | -1.852 | 0.542 | 0.0006 | 0.157 | 0.052 – 0.439 | -1.341 | 0.554 | 0.0155 | 0.262 | 0.084 – 0.756 |

Nagelkerke $R^2 = 0.196$ (95% efficacy); 0.238 (99% efficacy).
Tjur's $R^2 = 0.152$ (95% efficacy); 0.145 (99% efficacy).

### Ivermectin treatment in goats experimentally infected with Haemonchus contortus

All raw data from experimental infections are provided in S2 Data. Ivermectin was found to be not fully effective against strongyle nematodes in all trials reported in the present study. Larval cultures post treatment detected only *H. contortus* type L3 after treatment while pre-treatment samples also contained *Trichostrongylus* spp. and Chabertiidae. In order to confirm resistance of *H. contortus* in the local GIN communities, *H. contortus* adult females were obtained from abomasa of goats at the abattoirs in Kass and Nyala and used to obtain monospecific local *H. contortus* isolates. These isolates were used to experimentally infect goats. The level of ivermectin resistance that was observed in naturally infected goats was significantly (p < 0.05) higher in Bulbul, Nyala Domaia and Nyala University farm than in Kass. These findings were confirmed using male goats that were experimentally infected with the local *H. contortus* isolates from Nyala and Kass. In these experiments, ivermectin efficacy was evaluated at different time points after oral or SC injection as summarized in Table 6 and Fig 3. Remarkably, reduced ivermectin efficacy was evident in *H. contortus* isolates from Nyala independently of the route of administration (oral or SC injection) or of the day used for resampling after treatment.

**Fig 3.**
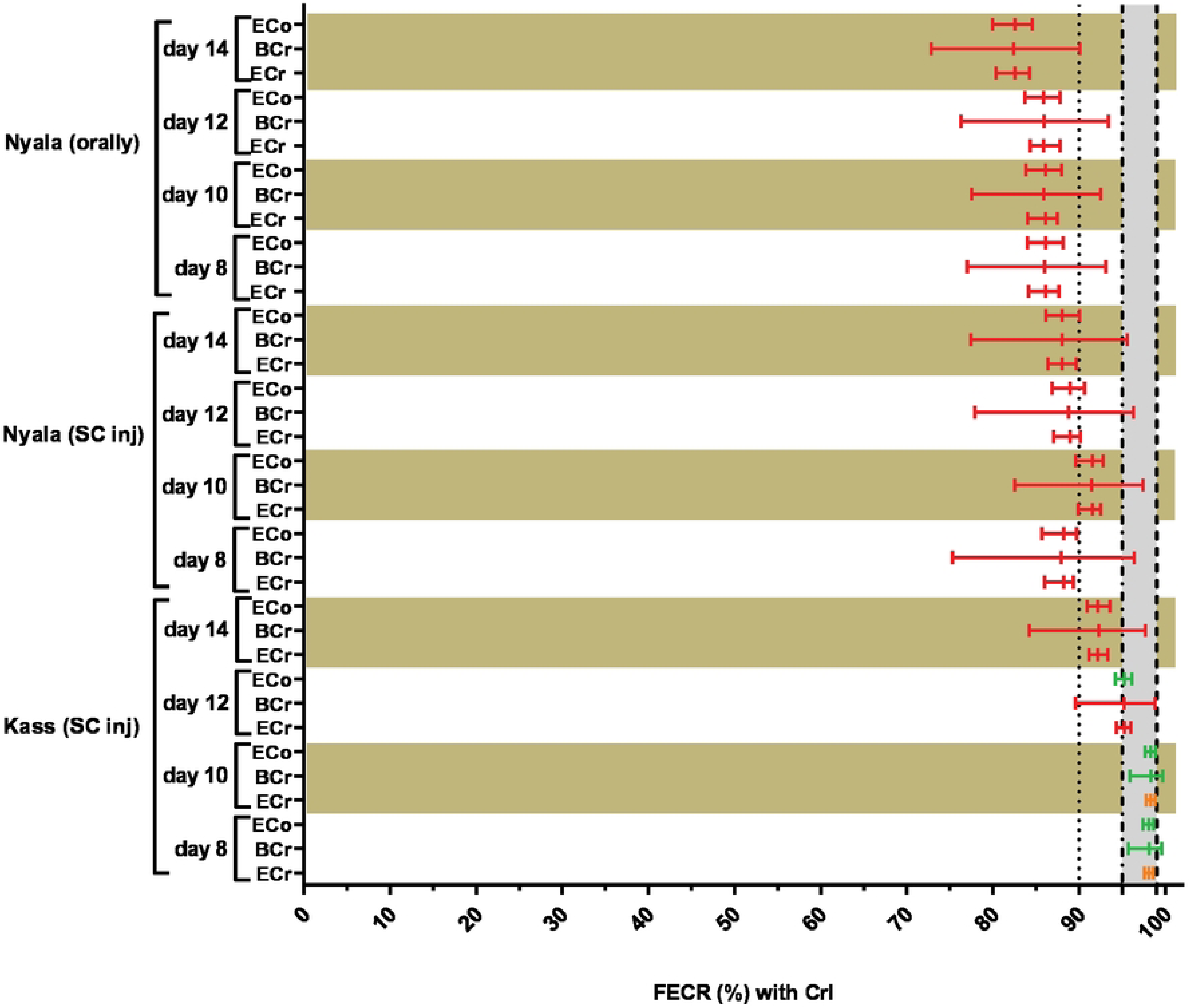
Faecal egg count reductions (FECRs) with both 90% or 95% credible intervals (CrIs) of male goats experimentally infected with *Haemonchus contortus* isolates collected from local abattoirs at Kass and Nyala, South Darfur, Sudan, post either subcutaneous or oral administration of 0.4 mg/kg body weight ivermectin on day 25 of infection, using different statistical analyses methods. Data were analysed separately for different days post treatment. ECr: paired FECRs and 90% credible intervals (CrIs) were calculated in R 4.1.1 using the eggCounts Bayesian hierarchical model version 2.3-2 without zero-inflation option and without individual efficacy. BCr: the 90% CrIs of the FECR for paired data were calculated using the Bayesian-Frequentist Hybrid Inference method through an open-source webpage using the R package bayescount version 1.1.0 (https://www.fecrt.com). The average reduction in FECs after treatment was calculated separately. ECo; the paired FECRs and 95% CrIs were calculated in R 4.1.1 using the eggCounts Bayesian hierarchical model version 2.3-2 without zero-inflation option and without individual efficacy. Results for ECr and BCr were assigned to the status resistance (red colour), low resistance (orange), susceptible (green) or inconclusive (black) as recommended in the revised guideline [25]. Results of the ECo were assigned resistance (red colour), susceptible (green) or inconclusive (black) as in the original guideline [34]. The grey zone indicates the range between the lower efficacy target of 95% and the expected efficacy of 99%.

**Table 6.** Faecal egg count reduction (FECR) (and 90% credible interval (CrI)) with male goats experimentally infected with *Haemonchus contortus* isolates collected from local abattoirs of Kass and Nyala, South Darfur, Sudan, before and after either subcutaneous or oral administration of 0.4 mg/kg body weight ivermectin on day 25 of infection to the treated groups.

| Source of infected materials | Route of administration | No. of animals included in the test<sup>a</sup> | Day of sample collection | Total eggs counted (Pre-treatment) | Mean faecal egg counts (eggs per gram) |  | FECR<sup>b</sup> (%) (90% CrI) | Test outcome<sup>c</sup> | Required sample size<sup>d</sup> |
| --- | --- | --- | --- | --- | --- | --- | --- | --- | --- |
|  |  |  |  |  | Pre-treatment | Day 12 |  |  |  |
| Kass | Subcutaneous injection | 10 | 8 | 1916 | 1916 | 36 | 98.1 (97.6 – 98.6) | Low resistant | 5 |
|  |  |  | 10 | 1916 | 1916 | 32 | 98.3 (97.8 – 98.8) | Low resistant |  |
|  |  |  | 12 | 1916 | 1916 | 92 | 95.3 (94.3 – 96.0) | Resistant |  |
|  |  |  | 14 | 1916 | 1916 | 148 | 92.2 (91.1 – 93.3) | Resistant |  |
| Nyala | Subcutaneous injection | 10 | 8 | 1386 | 1386 | 168 | 88.2 (86.0 – 89.4) | Resistant | 5 |
|  |  |  | 10 | 1386 | 1386 | 120 | 91.5 (89.9 – 92.5) | Resistant |  |
|  |  |  | 12 | 1386 | 1386 | 156 | 88.9 (87.0 – 90.2) | Resistant |  |
|  |  |  | 14 | 1386 | 1386 | 166 | 88.0 (86.4 – 89.7) | Resistant |  |
|  | Orally | 10 | 8 | 1430 | 1430 | 200 | 86.1 (84.2 – 87.6) | Resistant | 5 |
|  |  |  | 10 | 1430 | 1430 | 202 | 86.1 (84.0 – 87.5) | Resistant |  |
|  |  |  | 12 | 1430 | 1430 | 201 | 85.8 (84.3 – 87.8) | Resistant |  |
|  |  |  | 14 | 1430 | 1430 | 252 | 82.5 (80.3 – 84.2) | Resistant |  |

<sup>a</sup>Number of treated animals.
<sup>b</sup>FECRs were paired calculated by comparing data post and pre-treatment without zero-inflation option and without individual efficacy.
<sup>c</sup>The outcome of FECRs were identified as resistant, low resistant, susceptible or inconclusive based on the revised WAAVP guideline [25].
<sup>d</sup>Required sample size for a faecal egg count reduction test as per the revised WAAVP guideline [25], calculated using open source bayescount package version 1.1.0 (https://www.fecrt.com).

### Ivermectin treatment in sheep experimentally infected with nematodes surviving ivermectin treatment in goats

To exclude limited bioavailability of ivermectin as a factor causing insufficient efficacy of ivermectin in goats, nematodes surviving ivermectin treatment in goats in Bulbul and Nyala were used to obtain infective L3, which were then used to experimentally infect male sheep. Experimentally infected sheep received increasing doses of ivermectin every eight days starting with the dose recommended for sheep (0.2 mg/kg bw) and doubling this dose every week. After an initial stepwise decrease from a mean of 5133 before treatment to 730 on day 32, when the highest dose of 1.6 mg/kg bw was administered, egg counts started to rise again stepwise. The higher dosage had only small effects on FECs, and after applying the highest dose the FEC even increased (Table 7; Fig 4). For none of the comparisons, a FECR below 86.0% was observed confirming that these parasites also have a highly resistant phenotype when infecting sheep. At necropsy, only adult male and female *H. contortus* were isolated from the gastrointestinal tract of the infected sheep.

**Fig 4.**
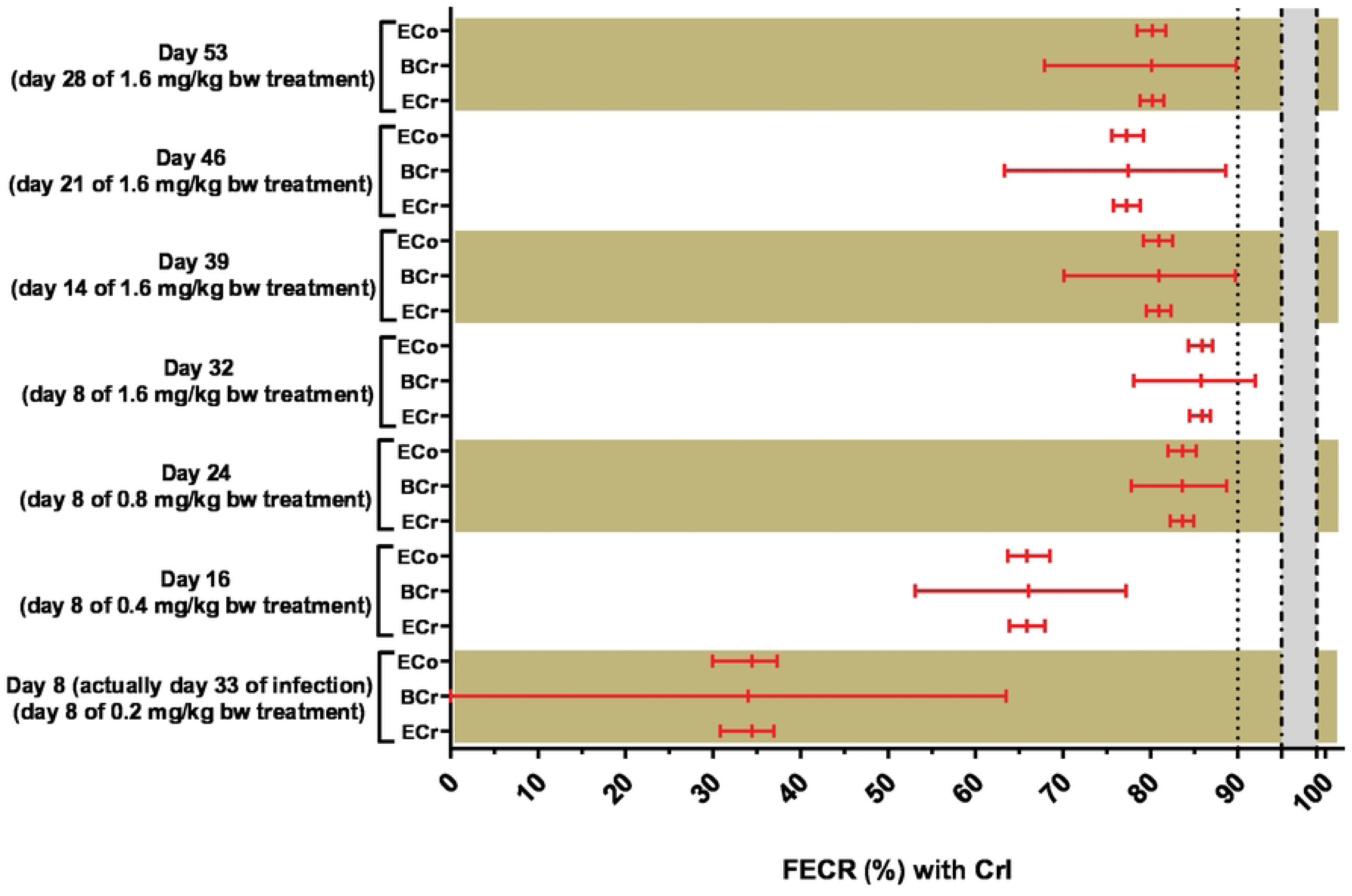
Comparisons between the faecal egg count reductions (FECRs) with both 90% and 95% credible intervals (CrIs) with male sheep (*n*=6) experimentally infected with nematodes surviving ivermectin treatment in goats from Bulbul and Nyala, South Darfur, Sudan, pre and post-subcutaneous administration of ivermectin with increasing doses, using different statistical analyses methods. Treatments with ivermectin were repeated every eight days with increasing doses, starting with the recommended sheep dose (0.2 mg/kg body weight) and ending at an eight-time higher dosage (1.6 mg/kg bw). Strongyle eggs were the only nematode eggs differentiated in faecal samples pre and post-treatment. ECr: paired faecal egg count reductions (FECRs) and 90% credible intervals (CrIs) were calculated in R 4.1.1 using the eggCounts Bayesian hierarchical model version 2.3-2 without zero-inflation option and without individual efficacy. BCr: the 90% CrIs of the FECR for paired data were calculated using the Bayesian-Frequentist Hybrid Inference method through an open-source webpage using the R package bayescount version 1.1.0 (https://www.fecrt.com). The average reduction in FECs after treatment was calculated separately. ECo; the paired FECRs and 95% CrIs were calculated in R 4.1.1 using the eggCounts Bayesian hierarchical model version 2.3-2 without zero-inflation option and without individual efficacy. Results for ECr and BCr were assigned to the status resistance (red colour), low resistance (orange), susceptible (green) or inconclusive (black) as recommended in the revised guideline [25]. Results of the ECo were assigned resistance (red colour), susceptible (green) or inconclusive (black) as in the original guideline [34]. The grey zone indicates the range between the lower efficacy target of 95% and the expected efficacy of 99%.

**Table 7.** Faecal egg count reduction (FECR) (and 90% credible interval (CrI)) with male sheep (*n*=6) experimentally infected with nematode surviving ivermectin treatment in goats from Bulbul and Nyala, South Darfur, Sudan, before and after subcutaneous administration of ivermectin with increasing doses.

| GI nematode | No. of animals included in the test<sup>b</sup> | Dose (mg/kg bw)<sup>c</sup> | Day of sample collection | Total eggs counted (Pre-treatment) | Mean faecal egg counts (eggs per gram) |  | FECR<sup>d</sup> (%) (90% CrI) | Test outcome<sup>e</sup> | Required sample size<sup>f</sup> |
| --- | --- | --- | --- | --- | --- | --- | --- | --- | --- |
|  |  |  |  |  | Pre-treatment | Day 12 |  |  |  |
| Strongyles<sup>a</sup> | 6 | 0.2 | 8 | 3080 | 5133 | 3387 | 34.5 (30.8 – 37.0) | Resistant | 6 |
|  |  | 0.4 | 16 | 3080 | 5133 | 1743 | 65.9 (63.9 – 67.9) | Resistant |  |
|  |  | 0.8 | 24 | 3080 | 5133 | 840 | 83.7 (82.3 – 84.9) | Resistant |  |
|  |  | 1.6 | 32 | 3080 | 5133 | 730 | 85.9 (84.5 – 86.8) | Resistant |  |
|  |  |  | 39 | 3080 | 5133 | 977 | 81.0 (79.5 – 82.4) | Resistant |  |
|  |  |  | 46 | 3080 | 5133 | 1157 | 77.3 (75.8 – 78.9) | Resistant |  |
|  |  |  | 53 | 3080 | 5133 | 1020 | 80.2 (78.8 – 81.5) | Resistant |  |

<sup>a</sup>Strongyle eggs were the only identified nematode eggs under the microscope in samples collected pre and post-treatment.
<sup>b</sup>Number of treated animals.
<sup>c</sup>Male sheep were received a first treatment with ivermectin at day 25 of infection.
<sup>d</sup>FECRs were paired calculated by comparing data post and pre-treatment without zero-inflation option and without individual efficacy.
<sup>e</sup>The outcome of FECRs were identified as resistant, low resistant, susceptible or inconclusive based on the revised WAAVP guideline [25].
<sup>f</sup>Required sample size for a faecal egg count reduction test as per the revised WAAVP guideline [25], calculated using open source bayescount package version 1.1.0 (https://www.fecrt.com).
*Abbreviations*; bw; body weight.

## Discussion

Ivermectin resistance is a severe problem regarding nematodes of ruminants in many geographic regions and the problem is considered a substantial threat for animal health and productivity worldwide [40]. While evidence of ivermectin resistance in sheep and goat parasites is widespread in commercial breeding systems [41], only a limited number of reports are available from Africa [42, 43]. However, in order to limit the spread of anthelmintic resistance, the extend of the problem needs to be precisely described to develop promising intervention strategies. In Sudan, particularly in South Darfur, the situation regarding resistance against benzimidazoles in cattle and goats has been characterised during the last years [2, 21]. The main goal of the present study was to evaluate the anthelmintic efficacy of ivermectin in sheep and goats naturally infected with GINs. Moreover, experimental infections were used to confirm that poor ivermectin efficacy is due to resistant *H. contortus* populations and that a higher metabolism in goats was not a reason for poor efficacy.

To the knowledge of the authors, this is the first comprehensive study to assess ivermectin efficacy in sheep and goats against GIN infections in Sudan. According to faecal cultures, animals in the trials involving naturally infected animals showed mixed infections before treatments, dominated by *Haemonchus* spp. but also including *Trichostrongylus* spp. and Chabertiidae. In contrast, only larvae of *Haemonchus* spp. were identified in faecal cultures after treatments.

Our data analysis of the sheep natural infection trials we performed showed the presence of ivermectin resistance in Nyala Domaia not only to the recommended dose but even to twice the recommended sheep dose. A therapeutic dose of ivermectin was insufficiently active (90 CrI: 73.6 – 77.5%) [25] against GINs. In 2018, a study by Alnaeim et al. [44] in sheep at the farm of the College of Veterinary Medicine, Sudan University of Science and Technology, Khartoum, Sudan, showed a lower efficacy to ivermectin (49.0% FECR) than our findings, but this study was not really representative due to the low number of only eight animals included. The efficacy reported here for sheep in South Darfur was lower than in Nejo District in Ethiopia (92.0% FECR) [45], but higher than the report from Gauteng Province in South Africa (67.0 – 87.0% FECR) [46] and in line with a previous report from Kajiado County in Kenya (76.0 – 86.0% FECR) [47].

In the present study, the efficacy of ivermectin was comprehensively studied in goats in the four different study locations using different treatment plans. Ivermectin at a therapeutic dose was ineffective in all four different study locations [25]. These findings on the occurrence of ivermectin resistance in goats in South Darfur corroborate reports from many regions of the world, and *H. contortus* is involved in most regions. In South America, the status of ivermectin resistance is critical, particularly in Brazil [48, 49]. Remarkably, in Europe occurrence of ivermectin resistance in goats has been described from regions such as Czech Republic, Germany, Romania and United Kingdom [12, 50–52]. Also, ivermectin resistance has been identified as problematic for small ruminant farming in Australia [41] and parts of Asia, including China [53], Malaysia [54] and Pakistan [55]. Noteworthy, pasture contamination with nematodes resistant to ivermectin has dramatically increased in Africa as many reports documented the development of resistance in the last decade, particularly for subsistence farming systems, including Ethiopia [13], Nigeria [15] and Uganda [16]. This might be considered as unexpected since subsistence farming would be expected to be associated with low treatment frequency and thus low selection pressure. Furthermore, despite the fact that ivermectin came on the market decades after the first benzimidazoles, resistance against ivermectin was as widespread as resistance against albendazole in goat parasites in the same regions [2]. One important aspect in this context in Sudan might be the fact that the subcutaneous and oral formulations of ivermectin are packaged in sizes allowing treatments of a flexible range of animal numbers starting from a single animal to a group of animals up to 100 heads. In contrast, package sizes for albendazole start with the smallest size sufficient to treat 50 sheep (Mohammedsalih, personal observation). This makes ivermectin cheaper than albendazole for farmers with only a very small number of animals. An additional advantage that makes ivermectin attractive is his activity against arthropods such as mites and lice [56].

Ivermectin was found to be effective against another identified nematode, *S. papillosus*, in goats in Bulbul and Nyala University farm. This is an interesting finding as the results of many studies from different regions supported our finding and detected no *S. papillosus* eggs in faeces of sheep and goats after ivermectin treatment [57–59].

In the present study, ivermectin FECR efficacy against GIN was also studied using twice the recommended sheep and goat doses. The efficacy of the higher dosage was significantly (p < 0.5) increased in both host species, but still the obtained values of the 90% CrIs were below the 99% threshold for resistance [25]. Noteworthy, strongyles from sheep in Nyala Domaia and goats in Bulbul were assigned resistant to ivermectin even at the dose increased twice the recommended to any animal-species. Lloberas et al. [60] observed 0% efficacy to ivermectin at a double dose in sheep experimentally infected with *H. contortus* isolates highly resistant to ivermectin. Our finding also agrees with a previous study from New Zealand reporting a calculated 62.0% efficacy to a double ivermectin dose in sheep experimentally infected with an ivermectin-resistant *Trichostrongylus colubriformis* isolate [61]. Increasing ivermectin doses accounted for an enhancement of drug exposure in the target tissues, as well as higher drug concentrations within the infected parasites [60]. Although, this change has the potential to kill weakly ivermectin resistant worms, it is obviously not sufficient to eliminate the complete population of resistant *H. contortus* present in South Darfur. In the long term it even comes with the risk to accelerate the selection of resistance through the increase of pasture contamination with highly resistant genotypes [62].

A third treatment plan was tested in the present study, in which goats in Bulbul and Nyala University farm were first treated with a therapeutic dose and those animals that continued shedding more than 150 strongyle EPG were retreated with 0.8 mg/kg bw dose on day 12. This plan was adapted from the farmer’s local habit in South Darfur, as they often retreat animals with anthelmintics after two weeks [2]. In contrast to the direct application of twice the recommended goat dose in the first treatment, using the higher dose in a second treatment even significantly decreased the efficacy significantly. This can be easily explained by removal of the ivermectin susceptible specimen in the strongyle communities by the first treatment leaving only ivermectin resistant nematodes in the animals when retreatment was performed.

In naturally infected sheep and goats, only *Haemonchus* spp. were detected in larval cultures after treatment. *Haemonchus contortus* in small ruminants is ranked first among different ruminant nematodes to evolve resistance against the available anthelmintics [3, 63], which has been attributed to its very high reproductive potential which resulted in huge population sizes [64]. The findings from natural infections in sheep and goats were confirmed in experimental infections using L3 cultured from eggs extracted from mature gravid female *H. contortus* goats from Kass and Nyala and used to infect worm-free goats. Irrespectively of the application route (oral vs. SC), ivermectin resistance was evident in both *H. contortus* isolates according to the revised WAAVP guideline FECR [25]. Similar findings were shown by Lespine et al. [17] as goats experimentally infected with GINs showed no significant differences in ivermectin efficacy between the oral and SC injection. For the Kass isolate, the analysis of data from early timepoints after treatment characterized the isolate as having a low resistance phenotype while full resistance was only detectable on days 12 and 14 post-treatment. This finding is in agreement with our previous study in goats using albendazole [2], and in line with recommendation of the revised WAAVP guideline [25]. Unexpectedly, however, efficacies observed in the experimental trials were significantly higher than those observed for naturally infected goats from the same region although the latter were infected with a mix of species that included apparently susceptible *Trichostrongylus* spp. and Chabertiidae. Possible explanations for this effect could be that the *Haemonchus* spp. from the naturally infected goats came from more sources and might therefore represent a mixture of different resistance pheno-and genotype, while the isolates were obtained from a limited number of goats and by chance might have represented only a subset of the population. An alternative explanation would be that the health status of the experimentally infected goats was in general better than that of the naturally infected animals and this can well have effects on the observed FECR [65].

Studying ivermectin efficacy in sheep experimentally infected with nematodes surviving ivermectin treatment in goats from Bulbul and Nyala further enhanced the power of the findings of the study. Since goats have a higher xenobiotic metabolism than sheep and most drugs have not been properly licensed for goats, there is a rule of thumb to use 1.5 time (levamisole) to two times the sheep dose (most other anthelmintics) [25], However, in the absence of data from studies conducted to licence a drug in a certain host species, the value of 99% for the expected efficacy is only poorly substantiated with data. This problem was circumvented by infecting sheep with parasites that survived treatment in goats and then testing this parasite population in a host species for which the anthelmintic is fully licensed. The fact that the offspring of *H. contortus* that survived ivermectin treatment in goats were also able to survive treatment in sheep finally confirms the resistance status level. By confirming the ivermectin resistance status, this trial excluded limited bioavailability of ivermectin as a confounder causing insufficient efficacy of this drug in goats. During inspection of the gastrointestinal tract of the experimentally infected sheep only adult *H. contortus* were identified, further confirming that this is the species that evolved ivermectin resistance in South Darfur.

The FECR was calculated from faecal egg counts pre and post-treatment using two statistical packages, both using Bayesian approaches, with interpretation based on the revised guideline [25]. Despite significant and considerable differences in the width of the 90% CrI, both statistical packages assigned the same status to all naturally infected groups (Cohen’s κ = 1). This is well in line to a previous study, in which these two statistical packages were compared regarding the assignment of a resistance status according to the revised guideline [66]. In this study, minimal differences in the assignment were observed, leading to a Cohen’s κ of 0.774. In contrast, comparison of the original and the revised guideline based on the same software package (eggCounts) revealed considerable differences (Cohen’s κ = 0.444). This observation that the guideline used for interpretation is of much higher importance than the statistical package used to analyse the data is of very high importance. First, it means that it will be almost impossible to compare results from future analyses to analyses in the past if not either the raw data are available in repositories, which is rarely the case for historical data, or 95% CIs or CrI are provided in new studies although they are no longer relevant in the assignment of the data. Even if the latter is the case, the number of methods used to calculate 95% CIs (and CrIs) has increased a lot over time and thus, it will be almost impossible to compare FECR data over time. This is of course a major problem since it prevents us from judging whether the incidence of anthelmintic resistance is increasing over time. Secondly, it also suggests that the revised guideline is pretty robust regarding different (at least Bayesian) statistical approaches and that different methods only lead to different results if parasite communities have a borderline status between susceptible and resistant [66].

In South Darfur, farmers are often treating diseased sheep and goats without prior veterinary examination, depending on their self-experience in diagnosis and selection of drugs. However, the treatment strategy of these farmers is usually depended on the use of oxytetracycline and ivermectin, since they have anecdotal evidence that the two drugs have had the potential to cure most animal diseases (Mohammedsalih, personal observations). This local habit increases the frequency of ivermectin treatments which plays a major role regarding the selection of resistance. Noteworthy, under-dosing is one of the major factors accelerating anthelmintic resistance evolution [67], and was found to occur due to many reasons including mistakes in dose calculation or human errors involving dose application. In Sudan, the recommended therapeutic dose of ivermectin (and other anthelmintics) is identical for sheep and goats (0.2 mg ivermectin/kg bw) for most package leaflets in veterinary pharmacies (Mohammedsalih, personal observation) indicating regular under-dosing in goats. The bioavailability of ivermectin is remarkably different in goats compared to sheep, therefore goats are require (1.5 to) 2-fold the sheep recommended dose to achieve the same pharmacological effect [18, 19]. Farmers in South Darfur are practicing a constant dosing for adults and half the adult dose for treatments of young-animals (Mohammedsalih, personal observation). This treatment strategy does not take into account variations in the animal’s body weight at the same animal species and usually results in under-dosing. One additional factor that may potentially contribute to the selection of ivermectin resistance by *H. controtus* populations in South Darfur is the presence of poor quality anthelmintics in the veterinary market of Sudan. This factor was explored in two previous studies from Sudan which measured the actual concentrations of some anthelmintics, and calculated lower concentration of albendazole in an oral suspension than what the manufacturer claimed [68, 69]. To exclude this critical factor and avoid to observe pseudo-resistance due to inadequate design of therapeutic treatment trials, all anthelmintics used in the present study were imported from Germany, the Netherlands or the UK.

Presence of resistant nematode populations in South Darfur has become a major problem in veterinary medicine and can be at least partly attributed to the complexity of sheep and goat production systems. There is an urgent need to adopt appropriate strategies for parasite control, such as local husbandry and management systems, optimize deworming strategies, in addition to regular monitoring of anthelmintic efficacies. Targeted selective treatment approaches would be advisable in South Darfur, since this approach can reduce amount of anthelmintics used, therefore decrease selection of resistant genotypes [70], as well as reducing the costs for anthelmintic treatments. Since there is a lack of information and instructions on the appropriate use of ivermectin in Sudan, particularly in goats, the results of the present study could be important to promote optimised strategies for the control of infections with GIN in sheep and goats. The next step in South Darfur would include evaluation of combination treatments using two or more anthelmintics, which may have significant roles to overcome the emerging of resistance development.

## Conclusions

Ivermectin resistance in GINs is an obviously already widespread occurring and further emerging threat for animal health and production efficacy of sheep and goats in Sudan. Reduced ivermectin efficacy was observed in strongyle nematodes in sheep and goats in all four different study locations in South Darfur, i.e. Bulbul, Kass, Nyala Domaia and Nyala University farm, included in the study. The results were confirmed in experimentally infected sheep and goats using nematodes surviving ivermectin treatment in goats or L3 cultivated from eggs obtained from gravid female *H. contortus* from goats at Kass and Nyala. Routine studies to evaluate the efficacy of anthelmintic treatments as well as implementation of targeted selective treatment approaches in South Darfur are urgently needed to overcome the evolution and spread of anthelmintic resistance in subsistence farming systems.

## Supporting information

**S1 Data. Raw data from naturally infected animals.** (xlsx)

**S2 Data. Raw data from experimentally infected animals.** (xlsx)

## Acknowledgements

We are grateful to sheep and goat producers for allowing them to work on their farm with their animals. We also thank dean and staff of the Faculty of Veterinary Science, University of Nyala, for their kindness and allowing us to work on the farm and give us a space to conduct the experimental infection trials.

## Author Contributions

**Conceptualization:** Khalid M. Mohammedsalih, Gerald Coles, Georg von Samson-Himmelstjerna, Jürgen Krücken.

**Data curation:** Khalid M. Mohammedsalih, Gerald Coles, Georg von Samson-Himmelstjerna, Jürgen Krücken.

**Formal analysis:** Khalid M. Mohammedsalih, Jürgen Krücken.

**Funding acquisition:** Khalid M. Mohammedsalih; Georg von Samson-Himmelstjerna

**Investigation:** Khalid M. Mohammedsalih, Abdoelnaim I. Y. Ibrahim, Fathel-Rahman Juma, Abdalhakaim A. H. Abdalmalaik, Ahmed Bashar

**Project administration:** Khalid M. Mohammedsalih, Jürgen Krücken, Georg von Samson-Himmelstjerna

**Supervision:** Khalid M. Mohammedsalih, Ahmed Bashar, Gerald Coles, Georg von Samson-Himmelstjerna, Jürgen Krücken.

**Validation:** Khalid M. Mohammedsalih, Ahmed Bashar, Gerald Coles, Georg von Samson-Himmelstjerna, Jürgen Krücken.

**Visualization:** Khalid M. Mohammedsalih, Jürgen Krücken.

**Writing – original draft:** Khalid M. Mohammedsalih, Jürgen Krücken, Georg von Samson-Himmelstjerna.

**Writing – review & editing:** Khalid M. Mohammedsalih, Abdoelnaim I. Y. Ibrahim, Fathel-Rahman Juma, Abdalhakaim A. H. Abdalmalaik, Ahmed Bashar, Gerald Coles, Jürgen Krücken, Georg von Samson-Himmelstjerna.

## Consent for publication

Not applicable.

## Data availability statement

All relevant data are within the manuscript and its Supporting Information files.

## Competing interests

The authors declare no conflict of interest.

## Funding

The field work in Sudan was funded by the International Foundation for Science (IFS; www.ifs.se), Sweden, as a co-fund with the Organisation of Islamic Conference Standing Committee on Scientific and Technological Cooperation (COMSTECH; www.comstech.org) (Grant No. B/5806-2) awarded to Dr Khalid M. Mohammedsalih. This study was also supported by a grant for Young Scientist, Ministry of Higher Education and Scientific Research (www.mohe.gov.sd), Sudan (Voucher Number: RPNUSNP/0130/2020) awarded to Dr Khalid M. Mohammedsalih. Part of anthelmintics used in this study was obtained using internal funds of the FU Berlin (https://www.vetmed.fu-berlin.de/einrichtungen/zfi/we13/index.html). The funders had no role in study design, data collection and analysis, decision to publish, or preparation of the manuscript.

## Notes

### Competing Interest Statement

The authors have declared no competing interest.

